# microbiomedataset: A tidyverse-style framework for organizing and processing microbiome data

**DOI:** 10.1101/2023.09.17.558096

**Authors:** Xiaotao Shen, Michael P. Snyder

## Abstract

Microbial communities exert a substantial influence on human health and have been unequivocally associated with a spectrum of human maladies, encompassing conditions such as anxiety1, depression2, hypertension3, cardiovascular diseases4, obesity4,5, diabetes6, inflammatory bowel disease7, and cancer8,9. This intricate interplay between microbiota community structures and host pathophysiology has kindled substantial interest and spurred active research endeavors across various scientific domains. Despite significant strides in sequencing technologies, which have unveiled the vast diversity of microbial populations across diverse ecosystems, the analysis of microbiome data remains a formidable challenge. The complexity inherent in such data, compounded by the absence of standardized data processing and analysis workflows, continues to pose substantial hurdles. The tidyverse paradigm, comprised of a suite of R packages meticulously crafted to facilitate efficient data manipulation and visualization, has garnered considerable acclaim within the data science community10. Its appeal stems from its innate simplicity and efficacy in organizing and processing data10. In recent times, a plethora of tools have been devised to address distinct omics data processing and analysis needs, including notable initiatives such as the tidymass project11, tidyomics project12, tidymicro13, and MicrobiotaProcess13,14. However, a conspicuous gap persists in the form of a standardized, tidyverse-based package for seamless and rigorous microbiome data processing and analysis.

To address this burgeoning demand for standardized and reproducible microbiome data analysis, we introduce microbiomedataset, an R package that embraces the tidyverse ethos to furnish a structured framework for the organization and processing of microbiome data. Microbiomedataset offers a comprehensive, customizable solution for the management, structuring, and processing of microbiome data. Importantly, this package seamlessly integrates with established bioinformatics tools, facilitating its incorporation into existing analytical pipelines11,13,14,15. Within this manuscript, we proffer an in-depth overview of the microbiomedataset package, elucidating its multifarious functionalities. Moreover, we substantiate its utility through illustrative case studies employing a publicly available microbiome dataset. It is imperative to underscore that microbiomedataset constitutes an integral component of the larger tidymicrobiome project, accessible via www.tidymicrobiome.org. Tidymicrobiome epitomizes an ecosystem of R packages that share a coherent design philosophy, grammar, and data structure, collectively engendering a robust, reproducible, and object-oriented computational framework. This project's development has been guided by several key tenets: (1) Cross-platform compatibility, (2) Uniformity, shareability, traceability, and reproducibility, and (3) Flexibility and extensibility. We further expound upon the advantages inherent in adopting a tidyverse-style framework for microbiome data analysis, underscoring the pronounced benefits in terms of standardization and reproducibility that microbiomedataset offers. In sum, microbiomedataset furnishes an accessible and efficient avenue for microbiome data analysis, catering to both neophyte and seasoned R users alike.

## Introduction

Microbial communities exert a substantial influence on human health and have been unequivocally associated with a spectrum of human maladies, encompassing conditions such as anxiety^1^, depression^2^, hypertension^3^, cardiovascular diseases^4^, obesity^4,5^, diabetes^6^, inflammatory bowel disease^7^, and cancer^8,9^. This intricate interplay between microbiota community structures and host pathophysiology has kindled substantial interest and spurred active research endeavors across various scientific domains. Despite significant strides in sequencing technologies, which have unveiled the vast diversity of microbial populations across diverse ecosystems, the analysis of microbiome data remains a formidable challenge. The complexity inherent in such data, compounded by the absence of standardized data processing and analysis workflows, continues to pose substantial hurdles. The tidyverse paradigm, comprised of a suite of R packages meticulously crafted to facilitate efficient data manipulation and visualization, has garnered considerable acclaim within the data science community^10^. Its appeal stems from its innate simplicity and efficacy in organizing and processing data^10^. In recent times, a plethora of tools have been devised to address distinct omics data processing and analysis needs, including notable initiatives such as the tidymass project^11^, tidyomics project^12^, tidymicro^13^, and MicrobiotaProcess^13,14^. However, a conspicuous gap persists in the form of a standardized, tidyverse-based package for seamless and rigorous microbiome data processing and analysis.

To address this burgeoning demand for standardized and reproducible microbiome data analysis, we introduce microbiomedataset, an R package that embraces the tidyverse ethos to furnish a structured framework for the organization and processing of microbiome data. Microbiomedataset offers a comprehensive, customizable solution for the management, structuring, and processing of microbiome data. Importantly, this package seamlessly integrates with established bioinformatics tools, facilitating its incorporation into existing analytical pipelines^11,13,14,15^. Within this manuscript, we proffer an in-depth overview of the microbiomedataset package, elucidating its multifarious functionalities. Moreover, we substantiate its utility through illustrative case studies employing a publicly available microbiome dataset. It is imperative to underscore that microbiomedataset constitutes an integral component of the larger tidymicrobiome project, accessible via www.tidymicrobiome.org. Tidymicrobiome epitomizes an ecosystem of R packages that share a coherent design philosophy, grammar, and data structure, collectively engendering a robust, reproducible, and object-oriented computational framework. This project’s development has been guided by several key tenets: (1) Cross-platform compatibility, (2) Uniformity, shareability, traceability, and reproducibility, and (3) Flexibility and extensibility. We further expound upon the advantages inherent in adopting a tidyverse-style framework for microbiome data analysis, underscoring the pronounced benefits in terms of standardization and reproducibility that microbiomedataset offers. In sum, microbiomedataset furnishes an accessible and efficient avenue for microbiome data analysis, catering to both neophyte and seasoned R users alike.

## Result

### Implementation

The microbiomedataset package offers a meticulously standardized and reproducible workflow grounded in the principles of the tidyverse, designed to effectively manage and process microbiome datasets. This versatile tool seamlessly integrates with a multitude of bioinformatics resources and boasts compatibility with various file formats, facilitating its seamless integration into existing data workflows. Part of the dynamic tidymicrobiome project, an ongoing endeavor dedicated to curating and advancing microbiome data processing and analysis through the tidyverse philosophy, microbiomedataset provides a robust foundation for microbiome data handling. The package is readily accessible and supported on Mac OS, Windows, and Linux platforms, aligning with the ethos of open-source accessibility. For enhanced reliability and accessibility, we have deployed microbiomedataset on three distinct code hosting platforms: GitHub (https://github.com/tidymicrobiome/microbiomedataset), GitLab (https://gitlab.com/tidymicrobiome/microbiomedataset), and the official tidymicrobiome website (https://www.tidymicrobiome.org/). Any updates or modifications made to the package are synchronized across these platforms, ensuring that users can access and install microbiomedataset under varied internet connectivity scenarios. Central to the microbiomedataset ecosystem is the “microbiome_dataset” object, a standardized data structure meticulously designed to encompass a diverse array of microbiome data types, including expression datasets, sample information, variable details, and so on (**Fig. 1a**). This object distinguishes itself through its seamless conversion capability to external data structures from different tools^14,15,^ rendering microbiomedataset highly adaptable for diverse analytical needs (**Fig. 1b**).

**Figure 1.**
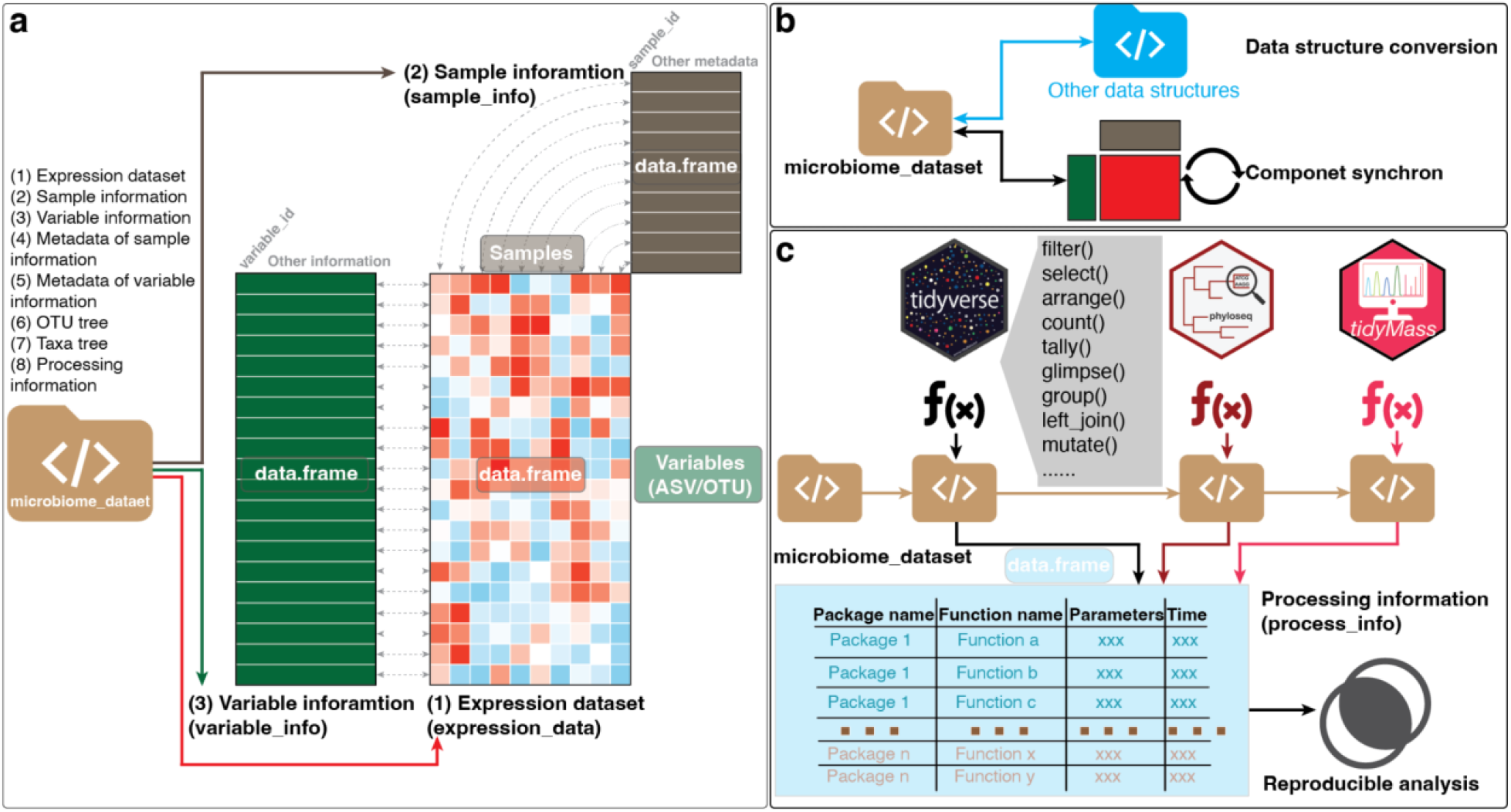
Microbiomedataset overview and its application in reproducible, object-oriented microbiome data processing and analysis. **(a)** The core of the microbiomedataset package is the “microbiome_dataset” object, a standardized data structure meticulously designed to accommodate diverse types of microbiome data, encompassing expression datasets, sample metadata, variable details, OTU (Operational Taxonomic Unit) trees, taxa trees, and pertinent processing information during the course of data analysis. This “microbiome_dataset” object serves as a unified data infrastructure tailored specifically for representing microbiome data. Notably, the tidymicrobiome project’s functions are explicitly compatible with this data format, with the added convenience of storing all requisite parameters within it. **(b)** The versatility of the “microbiome_dataset” object extends to its capacity for effortless conversion into alternative data structures. This adaptability enables users to seamlessly employ functions from the tidymicrobiome project or interface with external tools by transforming the “microbiome_dataset” object into their preferred data formats for subsequent processing and analysis. **(c)** The microbiomedataset package adheres to a modular and reproducible workflow rooted in the tidyverse style. Throughout the entire workflow, the uniform “microbiome_dataset” data structure remains pivotal. It consolidates all essential processing information and arguments, ensuring that the entirety of the analytical process remains transparent and readily reproducible.

Furthermore, any alterations made to a single dataset within the “microbiome_dataset” object result in the synchronized updating of all other associated data (**Fig. 1b**).

The development of microbiomedataset was underpinned by a modular architecture and an object-oriented workflow, both of which augment flexibility and extensibility, highly esteemed attributes in R package design. This object-oriented approach streamlines the data processing procedure while enhancing its transparency and intelligibility. The “microbiome_dataset” object maintains compatibility with most functions within the tidyverse, a potent tool for data manipulation in the R community. As an additional layer of safeguarding reproducibility, all processing steps and parameters are meticulously logged within the “microbiome_dataset” object (**Fig. 1c**).

### Applications

The workflow commences with the creation of the “microbiome_dataset” object, conveniently constructed using the “create_microbiome_dataset” function (**Fig. 2a**). Notably, microbiomedataset offers seamless compatibility with other data structures required by external tools (**Fig. 2a**). This flexibility allows users to effortlessly integrate microbiomedataset into their existing pipelines, including tools like phyloseq and MicrobiotaProcess^14^, enhancing its interoperability with established resources. Furthermore, users can easily extract specific components from the “microbiome_dataset” object. An essential aspect of the design is that the “microbiome_dataset” object seamlessly integrates with most functions from the tidyverse toolkit (**Fig. 2b**). This design choice obviates the need for users to acquaint themselves with novel functions, enabling them to employ familiar tools for microbiome data preprocessing tasks, such as sample and variable filtering, and data merging (**Fig. 2b**). Additionally, microbiomedataset empowers users with the ability to harness the ggplot2 data visualization toolkit for constructing visually appealing graphics in the ggplot2 style. This includes the creation of bar plots (**Fig. 2c**), circular plots (**Fig. 2d**), and phylogenetic trees (**Fig. 2e**) to enhance data visualization.

**Figure 2.**
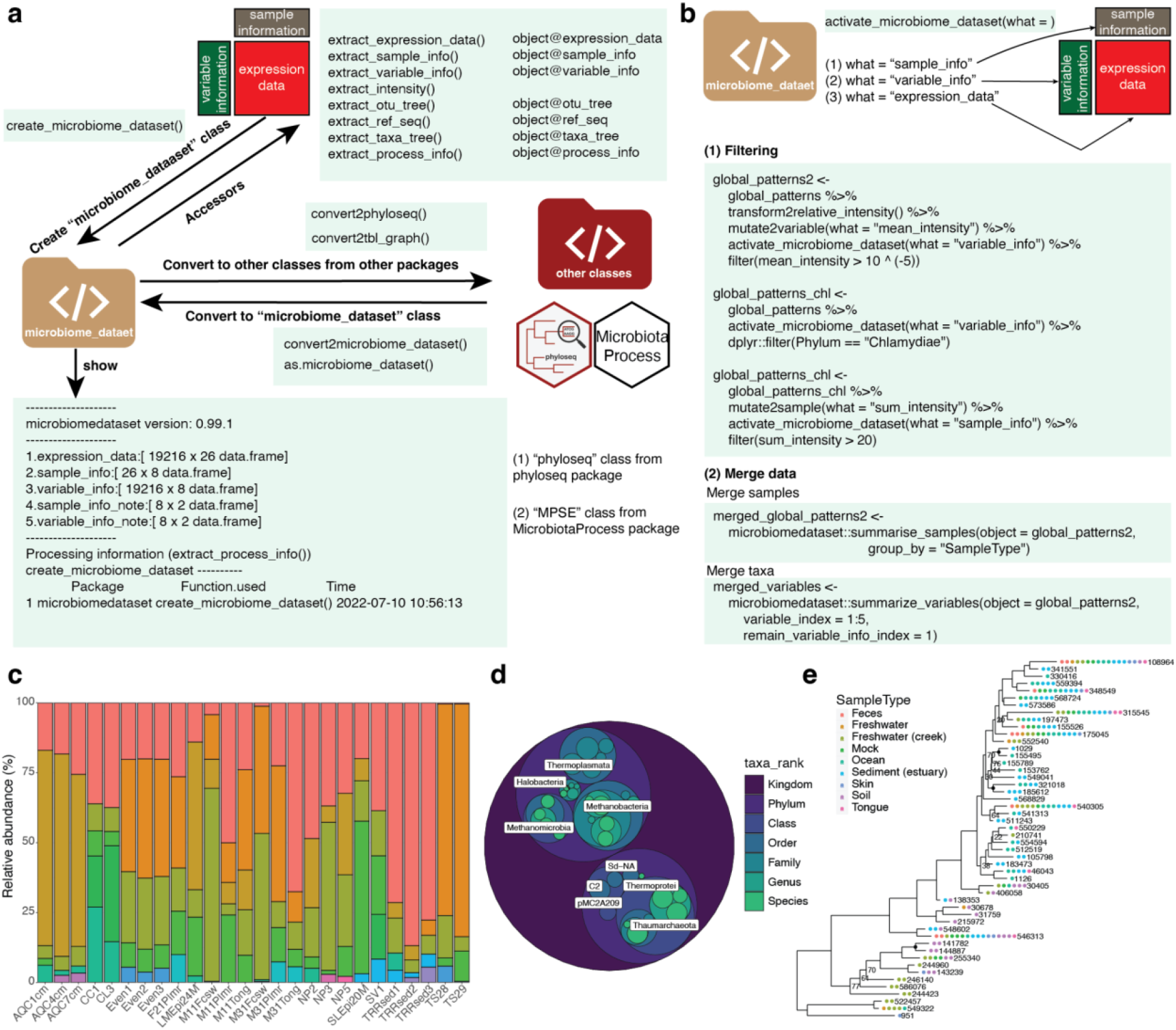
The microbiomedataset-based microbiome data processing and analysis pipeline. **(a)** The genesis of the “microbiome_dataset” object within the microbiomedataset package. Users can establish the “microbiome_dataset” object through the “create_microbiome_dataset” function. Alternatively, it allows for seamless conversion from other objects sourced from external tools, including but not limited to the phyloseq^15^ and MicrobiotaPorcess^14^ packages. Furthermore, the “microbiome_dataset” object can be flexibly transformed into various other data structures, and specific components can be extracted as needed. **(b)** Leveraging the power of the tidyverse-style functions for comprehensive data processing and analysis of microbiome datasets. A vast array of functions within the tidyverse toolkit can be seamlessly applied to the “microbiome_dataset” object, streamlining analytical workflows and ensuring compatibility with established data manipulation techniques. **(c)** Utilization of graphic functions from ggplot2 and other tools to facilitate data visualization for the “microbiome_dataset.” These visualization capabilities encompass a spectrum of formats, including bar plots **(c)**, circular plots **(d)**, and phylogenetic trees **(e)**, enhancing the accessibility and interpretability of microbiome data.

In a demonstrative case study, we applied microbiomedataset to process data from a published study originating from the phyloseq project^15^. Our results underscore the proficiency of microbiomedataset in organizing and processing microbiome data using standardized data structures and familiar tools like the tidyverse. These findings underscore microbiomedataset’s potential as a potent resource for conducting reproducible analyses of microbiome data (Supplementary Material).

## Discussion

Microbiome technologies have emerged as powerful tools for delving into the intricate relationships between microbial communities and their host organisms. Nevertheless, the field faces a persistent challenge in the form of data analysis, particularly due to the integration of diverse data types and the growing demand for personalized analytical approaches. In response to these challenges, the microbiomedataset and tidymicrobiome project (www.tidymicrobiome.org) provides a comprehensive solution through the introduction of the “microbiome_dataset” object (**Fig. 1a**). Built upon the foundational principles of tidyverse, microbiomedataset offers a suite of functions designed to unveil the characteristics and biological functions of microbial communities across varied environments. In summary, microbiomedataset confers significant advantages to the field of microbiome research, primarily in two pivotal areas. (1) Data sharing, tracing, and reproducible analyses. Microbiomedataset establishes a specific and standardized data structure within a holistic object-oriented workflow. This integrated framework streamlines data sharing, tracing, and reproducible analysis, enabling researchers to easily disseminate and replicate their analyses. Moreover, the uniform data structure facilitates the storage and tracking of parameters applied to metadata throughout the analysis pipeline, effectively addressing the longstanding challenge of inadequate metadata tracing in microbiome data analysis. (2) Flexibility and extensibility. Rooted in object-oriented and modular design principles, microbiomedataset seamlessly integrates with other tools utilized within the microbiome research community. This approach bestows upon microbiomedataset remarkable flexibility and extensibility, fostering collaboration and adaptability among researchers and tool developers in the dynamic field of microbiome.

To summarize, microbiomedataset is a flexible and comprehensive framework for organizing and processing microbiome data in a reproducible manner using the tidyverse style. It offers a standardized workflow and has been shown to be effective in the case study. Both novice and experienced R users can benefit from the package’s compatibility with other bioinformatics tools and customizable features. By promoting standardization and reproducibility in microbiome data analysis, we hope that microbiomedataset will improve the accuracy and understanding of microbiome research. The development team of the tidymicrobiome project (https://www.tidymicrobiome.org/) will provide long-term maintenance for the package.

## Author contributions

The method was conceived and its implementation was supervised by Xiaotao Shen. Xiaotao Shen was responsible for the development of the methods and packages, as well as the construction of the websites and the creation of help documents and tutorials. In addition, Xiaotao Shen prepared and conducted the analysis of the case study data and generated the figures. The manuscript was jointly authored by Xiaotao Shen and Michael Snyder. Chuchu Wang played a pivotal role in enhancing the quality of the manuscript. All authors contributed to the final version of the manuscript.

## Acknowledgments

We express our gratitude to Dr. Chuchu Wang for the invaluable guidance and insights provided during the figure preparation process. It is worth noting that this work did not receive any external funding.

## Conflict of Interest

Michael Snyder serves as a co-founder and holds membership on the scientific advisory boards of several entities, including Personalis, SensOmics, Filtricine, Qbio, January, Mirvie, and Oralome. Xiaotao Shen has no conflicts of interest to disclose.

## Data and code availability statement

Data sharing is not applicable to this article, as there were no new data generated or analyzed in the course of this study. For instructional purposes on how to utilize microbiomedataset, all demonstration data can be readily accessed on the tidymicrobiome website (https://www.tidymicrobiome.org/).

The microbiomedataset package, along with comprehensive documentation, is accessible via the official tidymicrobiome website: https://www.tidymicrobiome.org/. The complete source code for the microbiomedataset project is hosted on GitHub (https://github.com/microbiomedataset) and is made public under the MIT License. It is compatible with Windows, macOS X, and the majority of Linux distributions. Additionally, the source code for the case study is furnished as Supplementary material.

## Notes

https://www.tidymicrobiome.org/

## References

1. Yang, B., Wei, J., Ju, P. & Chen, J. Effects of regulating intestinal microbiota on anxiety symptoms: A systematic review. Gen Psychiatr 32, e100056 (2019).

2. Liu, L. et al. Gut microbiota and its metabolites in depression: from pathogenesis to treatment. EBioMedicine 90, 104527 (2023).

3. O’Donnell, J. A., Zheng, T., Meric, G. & Marques, F. Z. The gut microbiome and hypertension. Nat. Rev. Nephrol. 19, 153–167 (2023).

4. Rahman, M. M. et al. The Gut Microbiota (Microbiome) in Cardiovascular Disease and Its Therapeutic Regulation. Front. Cell. Infect. Microbiol. 12, 903570 (2022).

5. Van Hul, M. & Cani, P. D. The gut microbiota in obesity and weight management: microbes as friends or foe? Nat. Rev. Endocrinol. 19, 258–271 (2023).

6. Zhou, Z., Sun, B., Yu, D. & Zhu, C. Gut Microbiota: An Important Player in Type 2 Diabetes Mellitus. Front. Cell. Infect. Microbiol. 12, 834485 (2022).

7. Caruso, R., Lo, B. C. & Nú ñ ez, G. Host–microbiota interactions in inflammatory bowel disease. Nat. Rev. Immunol. 20, 411–426 (2020).

8. Clemente, J. C., Ursell, L. K., Parfrey, L. W. & Knight, R. The impact of the gut microbiota on human health: an integrative view. Cell 148, 1258–1270 (2012).

9. Afzaal, M. et al. Human gut microbiota in health and disease: Unveiling the relationship. Front. Microbiol. 13, 999001 (2022).

10. Wickham, H. & Grolemund, G. R for Data Science: Import, Tidy, Transform, Visualize, and Model Data. (‘O’Reilly Media, Inc.’, 2016).

11. Shen, X. et al. TidyMass an object-oriented reproducible analysis framework for LC-MS data. Nat. Commun. 13, 4365 (2022).

12. Hutchison, W. J. et al. The tidyomics ecosystem: Enhancing omic data analyses. bioRxiv 2023.09.10.557072 (2023) doi:10.1101/2023.09.10.557072.

13. Carpenter, C. M. et al. tidyMicro: a pipeline for microbiome data analysis and visualization using the tidyverse in R. BMC Bioinformatics 22, 1–13 (2021).

14. Xu, S. et al. A comprehensive R package for deep mining microbiome. Innovation (Camb) 4, 100388 (2023).

15. McMurdie, P. J. & Holmes, S. phyloseq: An R Package for Reproducible Interactive Analysis and Graphics of Microbiome Census Data. PLoS One 8, e61217 (2013).

