## Supplementary for "microbiomedataset: A tidyverse-style framework for organizing and processing microbiome data"

### 1. "microbiome_dataset" class

For one mass_dataset class object, we can get the summary information of it.

Data preparation.

```
library(microbiomedataset)
expression_data <-
  as.data.frame(matrix(
    sample(1:100, 100, replace = TRUE),
    nrow = 10,
    ncol = 10
  ))

rownames(expression_data) <- paste0("OTU", 1:nrow(expression_data))
colnames(expression_data) <-
  paste0("Sample", 1:ncol(expression_data))
expression_data
#>       Sample1 Sample2 Sample3 Sample4 Sample5 Sample6 Sample7 Sample8 Samp
le9
#> OTU1       78       9      91      67      47      28      58      57
 45
#> OTU2       34      96      94      40      61      62      42      33
 47
#> OTU3       74      47       2      36      70      75      77      42
 22
#> OTU4       98      63      43      94       3      71      51      89
 63
#> OTU5       77      81      31      92       7      78      19      73
 11
#> OTU6       80      76      38       8      48      79      50      28
 37
#> OTU7       61      39      32       9      72      59      22      19
 51
#> OTU8       75      98      13      63       4      80      65      81
  8
```

```
#> OTU9        26      67      93      64      40       8      42      14
 48
#> OTU10       28      22      26      68      14      22      63      45
 31
#>       Sample10
#> OTU1        78
#> OTU2        76
#> OTU3        84
#> OTU4        78
#> OTU5        14
#> OTU6        11
#> OTU7        62
#> OTU8        40
#> OTU9        34
#> OTU10       94

variable_info <-
  as.data.frame(matrix(
    sample(letters, 70, replace = TRUE),
    nrow = nrow(expression_data),
    ncol = 7
  ))

rownames(variable_info) <- rownames(expression_data)
colnames(variable_info) <-
  c("Domain",
    "Phylum",
    "Class",
    "Order",
    "Family",
    "Genus",
    "Species")

variable_info$variable_id <-
  rownames(expression_data)

sample_info <-
  data.frame(sample_id = colnames(expression_data),
             class = "Subject")

object <-
  create_microbiome_dataset(
    expression_data = expression_data,
    sample_info = sample_info,
    variable_info = variable_info
  )

object
#> --------------------
#> microbiomedataset version: 0.99.1
```

```
#> --------------------
#> 1.expression_data:[ 10 x 10 data.frame]
#> 2.sample_info:[ 10 x 2 data.frame]
#> 3.variable_info:[ 10 x 8 data.frame]
#> 4.sample_info_note:[ 2 x 2 data.frame]
#> 5.variable_info_note:[ 8 x 2 data.frame]
#> --------------------
#> Processing information (extract_process_info())
#> create_microbiome_dataset ----------
#>             Package              Function.used                Time
#> 1 microbiomedataset create_microbiome_dataset() 2023-04-19 17:01:20
```

### 2. Importing data

You can load the demo data in microbiomedataset package to see the microbiome_dataset class.

```
library(microbiomedataset)
library(tidyverse)

data("global_patterns")

global_patterns
#> --------------------
#> microbiomedataset version: 0.99.1
#> --------------------
#> 1.expression_data:[ 19216 x 26 data.frame]
#> 2.sample_info:[ 26 x 8 data.frame]
#> 3.variable_info:[ 19216 x 8 data.frame]
#> 4.sample_info_note:[ 8 x 2 data.frame]
#> 5.variable_info_note:[ 8 x 2 data.frame]
#> --------------------
#> Processing information (extract_process_info())
#> create_microbiome_dataset ----------
#>             Package              Function.used                Time
#> 1 microbiomedataset create_microbiome_dataset() 2022-07-10 10:56:13
```

So you can see that we have 1,9216 variables and 26 samples in the dataset.

Create microbiome_dataset class object.

You can also create the microbiome_dataset class using the create_microbiome_dataset function.

We need to prepare at least three pieces of data for it.

1. expression_data: rows are variables, and columns are samples.
2. sample_info: Information for all the samples in expression_data. The first column should be sample_id, which should be identical to the column names of expression_data.

3. variable_info: Information for all the variables in expression_data. The first column should be variable_id, which should be identical to the row names of expression_data.

```
expression_data <-
  as.data.frame(matrix(
    sample(1:100, 100, replace = TRUE),
    nrow = 10,
    ncol = 10
  ))

rownames(expression_data) <- paste0("OTU", 1:nrow(expression_data))
colnames(expression_data) <-
  paste0("Sample", 1:ncol(expression_data))

expression_data
#>       Sample1 Sample2 Sample3 Sample4 Sample5 Sample6 Sample7 Sample8 Samp
le9
#> OTU1       90      74      80      95      12      61      70      97
  6
#> OTU2       43      65      68      74      57      46      42      90
 73
#> OTU3       30      21      54      33      83       4      46       8
 51
#> OTU4       52      70      39      10      69      54      95      14
  1
#> OTU5       41      68      43      72      76      44      15      29
  4
#> OTU6       58      14      42      36       9      79      65       3
 94
#> OTU7       58      84       7       4      52      40      59      86
 46
#> OTU8       35      41      73      91      67      63      63      36
 28
#> OTU9       15       9     100      84      70      54      13      84
 31
#> OTU10      67      99      13      21      53      20      70      71
 13
#>       Sample10
#> OTU1        99
#> OTU2        52
#> OTU3        73
#> OTU4         5
#> OTU5        61
#> OTU6         2
#> OTU7        30
#> OTU8        26
#> OTU9        10
#> OTU10       59

variable_info <-
```

```r
  as.data.frame(matrix(
    sample(letters, 70, replace = TRUE),
    nrow = nrow(expression_data),
    ncol = 7
  ))

rownames(variable_info) <- rownames(expression_data)
colnames(variable_info) <-
  c("Domain",
    "Phylum",
    "Class",
    "Order",
    "Family",
    "Genus",
    "Species")

variable_info$variable_id <-
  rownames(expression_data)

variable_info <-
  variable_info %>%
  dplyr::select(variable_id, dplyr::everything())

sample_info <-
  data.frame(sample_id = colnames(expression_data),
             class = "Subject")

object <-
  create_microbiome_dataset(
    expression_data = expression_data,
    sample_info = sample_info,
    variable_info = variable_info
  )

object
#> --------------------
#> microbiomedataset version: 0.99.1
#> --------------------
#> 1.expression_data:[ 10 x 10 data.frame]
#> 2.sample_info:[ 10 x 2 data.frame]
#> 3.variable_info:[ 10 x 8 data.frame]
#> 4.sample_info_note:[ 2 x 2 data.frame]
#> 5.variable_info_note:[ 8 x 2 data.frame]
#> --------------------
#> Processing information (extract_process_info())
#> create_microbiome_dataset ----------
#>             Package               Function.used                Time
#> 1 microbiomedataset create_microbiome_dataset() 2023-04-19 17:02:31
```

Convert phyloseq class to microbiome_dataset class object.

We can also transfer or convert other common class objects to microbiome_dataset class.

Please install phyloseq package first.

```
if(!require(BiocManager)){
  install.packages("BiocManager")
}

if(!require(phyloseq)){
  BiocManager::install("phyloseq")
}

library(phyloseq)
data(GlobalPatterns)
GlobalPatterns
#> phyloseq-class experiment-level object
#> otu_table()   OTU Table:         [ 19216 taxa and 26 samples ]
#> sample_data() Sample Data:       [ 26 samples by 7 sample variables ]
#> tax_table()   Taxonomy Table:    [ 19216 taxa by 7 taxonomic ranks ]
#> phy_tree()    Phylogenetic Tree: [ 19216 tips and 19215 internal nodes ]
```

The first function is convert2microbiome_dataset:

```
object1 <-
  convert2microbiome_dataset(object = GlobalPatterns)
object1
#> --------------------
#> microbiomedataset version: 0.99.1
#> --------------------
#> 1.expression_data:[ 19216 x 26 data.frame]
#> 2.sample_info:[ 26 x 8 data.frame]
#> 3.variable_info:[ 19216 x 8 data.frame]
#> 4.sample_info_note:[ 8 x 2 data.frame]
#> 5.variable_info_note:[ 8 x 2 data.frame]
#> --------------------
#> Processing information (extract_process_info())
#> create_microbiome_dataset ----------
#>             Package                Function.used                Time
#> 1 microbiomedataset create_microbiome_dataset() 2023-04-19 17:02:42
```

The second function is as.microbiome_dataset:

```
object2 <-
  as.microbiome_dataset(object = GlobalPatterns)
object2
#> --------------------
#> microbiomedataset version: 0.99.1
#> --------------------
#> 1.expression_data:[ 19216 x 26 data.frame]
#> 2.sample_info:[ 26 x 8 data.frame]
```

```
#> 3.variable_info:[ 19216 x 8 data.frame]
#> 4.sample_info_note:[ 8 x 2 data.frame]
#> 5.variable_info_note:[ 8 x 2 data.frame]
#> --------------------
#> Processing information (extract_process_info())
#> create_microbiome_dataset ----------
#>             Package             Function.used                Time
#> 1 microbiomedataset create_microbiome_dataset() 2023-04-19 17:02:50

microbiomedataset::plot_barplot(object = object,
                                top_n = 5,
                                fill = "Phylum")
```

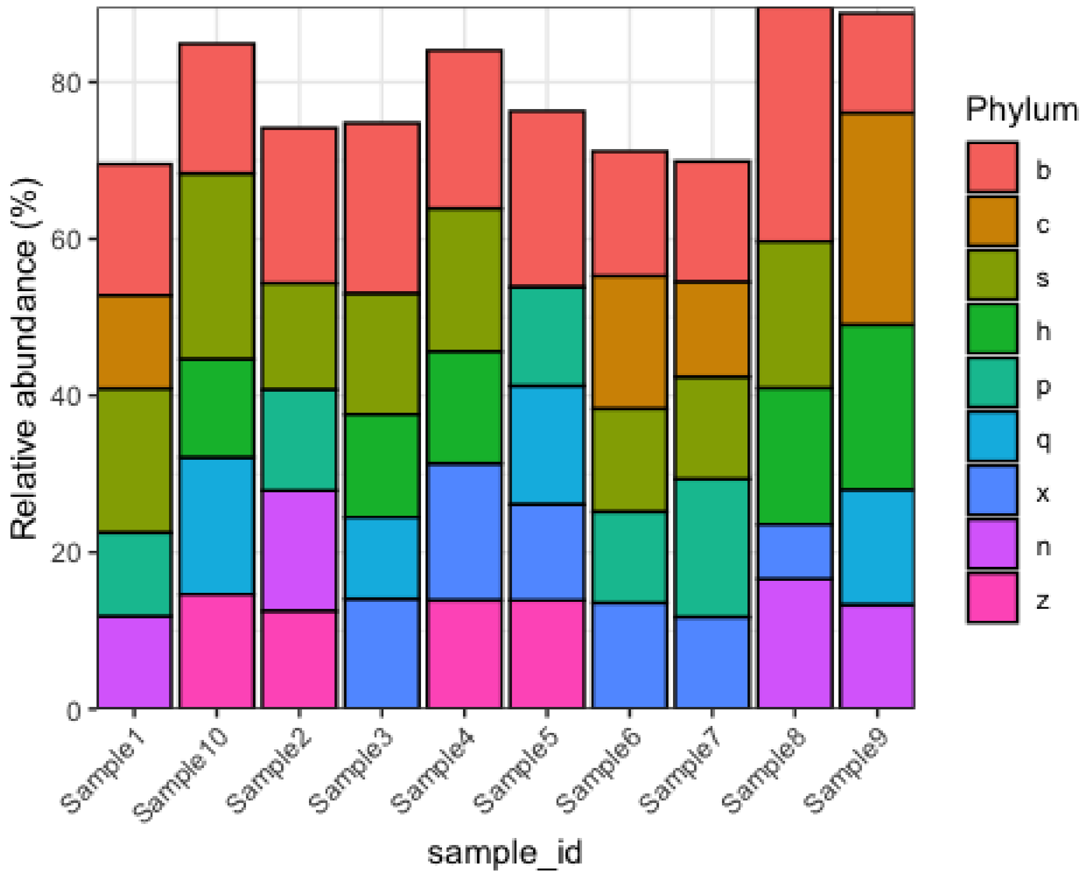


```
microbiomedataset::plot_barplot(object = object,
                                top_n = 5,
                                fill = "Phylum",
                                relative = TRUE,
                                re_calculate_relative = TRUE)
```

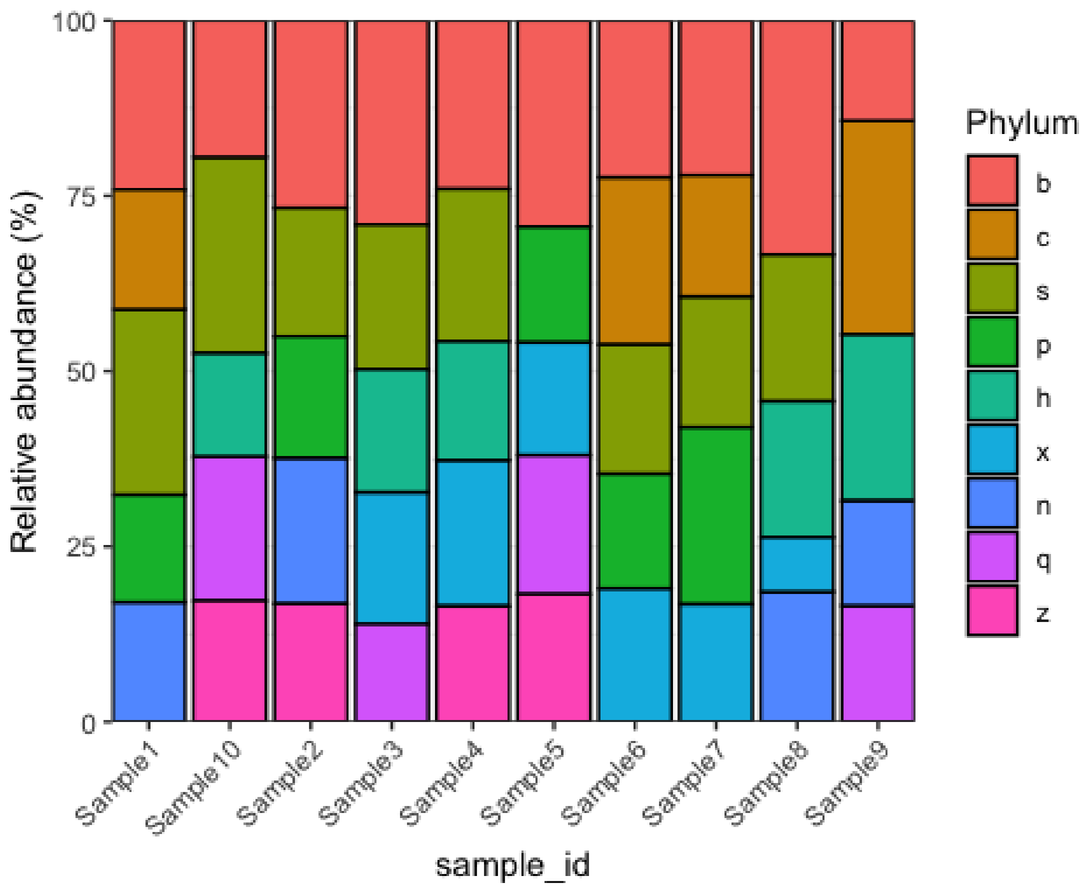


### 3. Transform absolute to relative intensity

Let’s load the demo data.

```
library(microbiomedataset)
library(tidyverse)
data("global_patterns")
global_patterns
#> --------------------
#> microbiomedataset version: 0.99.1
#> --------------------
#> 1.expression_data:[ 19216 x 26 data.frame]
#> 2.sample_info:[ 26 x 8 data.frame]
#> 3.variable_info:[ 19216 x 8 data.frame]
#> 4.sample_info_note:[ 8 x 2 data.frame]
#> 5.variable_info_note:[ 8 x 2 data.frame]
#> --------------------
#> Processing information (extract_process_info())
#> create_microbiome_dataset ----------
#>             Package             Function.used                Time
#> 1 microbiomedataset create_microbiome_dataset() 2022-07-10 10:56:13
```

```r
expression_data <-
  extract_expression_data(global_patterns)
expression_data$CL3 %>%
  head()
#> [1] 0 0 0 0 0 0

expression_data$CL3 %>%
  head()
#> [1] 0 0 0 0 0 0

apply(expression_data, 2, sum)
#>      CL3      CC1      SV1  M31Fcsw  M11Fcsw  M31Plmr  M11Plmr  F21Plmr
#>   864077  1135457   697509  1543451  2076476   718943   433894   186297
#>  M31Tong  M11Tong LMEpi24M SLEpi20M   AQC1cm   AQC4cm   AQC7cm      NP2
#>  2000402   100187  2117592  1217312  1167748  2357181  1699293   523634
#>      NP3      NP5  TRRsed1  TRRsed2  TRRsed3     TS28     TS29    Even1
#>  1478965  1652754    58688   493126   279704   937466  1211071  1216137
#>    Even2    Even3
#>   971073  1078241
```

The raw data is count.

```r
global_patterns_relative <-
  transform2relative_intensity(object = global_patterns)

expression_data_relative <-
  extract_expression_data(global_patterns_relative)
expression_data_relative$CL3 %>%
  head()
#> [1] 0 0 0 0 0 0

apply(expression_data_relative, 2, sum)
#>      CL3      CC1      SV1  M31Fcsw  M11Fcsw  M31Plmr  M11Plmr  F21Plmr
#>        1        1        1        1        1        1        1        1
#>  M31Tong  M11Tong LMEpi24M SLEpi20M   AQC1cm   AQC4cm   AQC7cm      NP2
#>        1        1        1        1        1        1        1        1
#>      NP3      NP5  TRRsed1  TRRsed2  TRRsed3     TS28     TS29    Even1
#>        1        1        1        1        1        1        1        1
#>    Even2    Even3
#>        1        1
```

### 4. Functions for accessing and preprocessing data

Let’s load the demo data.

```r
library(microbiomedataset)
library(tidyverse)
data("global_patterns")
global_patterns
#> --------------------
```

```
#> microbiomedataset version: 0.99.1
#> --------------------
#> 1.expression_data:[ 19216 x 26 data.frame]
#> 2.sample_info:[ 26 x 8 data.frame]
#> 3.variable_info:[ 19216 x 8 data.frame]
#> 4.sample_info_note:[ 8 x 2 data.frame]
#> 5.variable_info_note:[ 8 x 2 data.frame]
#> --------------------
#> Processing information (extract_process_info())
#> create_microbiome_dataset ----------
#>             Package               Function.used                Time
#> 1 microbiomedataset create_microbiome_dataset() 2022-07-10 10:56:13
```

#### 4.1 Accessors

```
dim(global_patterns)
#> variables   samples
#>     19216        26
nrow(global_patterns)
#> variables
#>     19216
ncol(global_patterns)
#> samples
#>      26

colnames(global_patterns)
#>  [1] "CL3"      "CC1"      "SV1"      "M31Fcsw"  "M11Fcsw"  "M31Plmr"
#>  [7] "M11Plmr"  "F21Plmr"  "M31Tong"  "M11Tong"  "LMEpi24M" "SLEpi20M"
#> [13] "AQC1cm"   "AQC4cm"   "AQC7cm"   "NP2"      "NP3"      "NP5"
#> [19] "TRRsed1"  "TRRsed2"  "TRRsed3"  "TS28"     "TS29"     "Even1"
#> [25] "Even2"    "Even3"

head(rownames(global_patterns))
#> [1] "549322" "522457" "951"    "244423" "586076" "246140"

extract_sample_info(global_patterns) %>%
  colnames()
#> [1] "sample_id"                 "Primer"
#> [3] "Final_Barcode"             "Barcode_truncated_plus_T"
#> [5] "Barcode_full_length"       "SampleType"
#> [7] "Description"               "class"

extract_variable_info(global_patterns) %>%
  colnames()
#> [1] "variable_id" "Kingdom"     "Phylum"      "Class"       "Order"
#> [6] "Family"      "Genus"       "Species"

extract_expression_data(global_patterns) %>%
  head()
#>        CL3 CC1 SV1 M31Fcsw M11Fcsw M31Plmr M11Plmr F21Plmr M31Tong M11Tong
#> 549322   0   0   0       0       0       0       0       0       0       0
```

```
#> 522457   0   0   0       0       0       0       0       0       0       0
#> 951      0   0   0       0       0       0       1       0       0       0
#> 244423   0   0   0       0       0       0       0       0       0       0
#> 586076   0   0   0       0       0       0       0       0       0       0
#> 246140   0   0   0       0       0       0       0       0       0       0
#>        LMEpi24M SLEpi20M AQC1cm AQC4cm AQC7cm NP2 NP3 NP5 TRRsed1 TRRsed2
#> 549322        0        1     27    100    130   1   0   0       0       0
#> 522457        0        0      0      2      6   0   0   0       0       0
#> 951           0        0      0      0      0   0   0   0       0       0
#> 244423        0        0      0     22     29   0   0   0       0       0
#> 586076        0        0      0      2      1   0   0   0       0       0
#> 246140        0        0      0      1      3   0   0   0       0       0
#>        TRRsed3 TS28 TS29 Even1 Even2 Even3
#> 549322       0    0    0     0     0     0
#> 522457       0    0    0     0     0     0
#> 951          0    0    0     0     0     0
#> 244423       0    0    0     0     0     0
#> 586076       0    0    0     0     0     0
#> 246140       0    0    0     0     0     0
```

```
extract_sample_info(global_patterns) %>%
  head()
#>   sample_id  Primer Final_Barcode Barcode_truncated_plus_T Barcode_full_le
ngth
#> 1       CL3 ILBC_01        AACGCA                   TGCGTT         CTAGCGT
GCGT
#> 2       CC1 ILBC_02        AACTCG                   CGAGTT         CATCGAC
GAGT
#> 3       SV1 ILBC_03        AACTGT                   ACAGTT         GTACGCA
CAGT
#> 4   M31Fcsw ILBC_04        AAGAGA                   TCTCTT         TCGACAT
CTCT
#> 5   M11Fcsw ILBC_05        AAGCTG                   CAGCTT         CGACTGC
AGCT
#> 6   M31Plmr ILBC_07        AATCGT                   ACGATT         CGAGTCA
CGAT
#>   SampleType                                Description   class
#> 1       Soil   Calhoun South Carolina Pine soil, pH 4.9 Subject
#> 2       Soil   Cedar Creek Minnesota, grassland, pH 6.1 Subject
#> 3       Soil Sevilleta new Mexico, desert scrub, pH 8.3 Subject
#> 4      Feces    M3, Day 1, fecal swab, whole body study Subject
#> 5      Feces   M1, Day 1, fecal swab, whole body study  Subject
#> 6       Skin    M3, Day 1, right palm, whole body study Subject
```

```
extract_variable_info(global_patterns) %>%
  head()
#>   variable_id Kingdom        Phylum        Class        Order         Famil
y
#> 1      549322 Archaea Crenarchaeota Thermoprotei         <NA>           <N
A>
```

```
#> 2      522457 Archaea Crenarchaeota Thermoprotei         <NA>          <N
A>
#> 3         951 Archaea Crenarchaeota Thermoprotei Sulfolobales Sulfolobacea
e
#> 4      244423 Archaea Crenarchaeota        Sd-NA         <NA>          <N
A>
#> 5      586076 Archaea Crenarchaeota        Sd-NA         <NA>          <N
A>
#> 6      246140 Archaea Crenarchaeota        Sd-NA         <NA>          <N
A>
#>        Genus                  Species
#> 1       <NA>                     <NA>
#> 2       <NA>                     <NA>
#> 3 Sulfolobus Sulfolobusacidocaldarius
#> 4       <NA>                     <NA>
#> 5       <NA>                     <NA>
#> 6       <NA>                     <NA>
```

#### 4.2 Preprocessing

The microbiomedataset package also includes functions for filtering, subsetting, and merging abundance data.

In the following example, the global_patterns data is first transformed to relative abundance, creating the new global_patterns2 object, which is then filtered such that only OTUs with a mean greater than 10^-5 are kept.

```
global_patterns2 <-
  global_patterns %>%
  transform2relative_intensity() %>%
  mutate2variable(what = "mean_intensity") %>%
  activate_microbiome_dataset(what = "variable_info") %>%
  filter(mean_intensity > 10 ^ (-5))
```

This results in a highly-subsetted object, global_patterns2, containing just 4624 of the original ~19216 OTUs.

Next, only remain the variables that phylum Chlamydiae.

```
global_patterns_chl <-
  global_patterns %>%
  activate_microbiome_dataset(what = "variable_info") %>%
  dplyr::filter(Phylum == "Chlamydiae")
```

Next, only remain the samples with total intensity > 20.

```
global_patterns_chl <-
  global_patterns_chl %>%
  mutate2sample(what = "sum_intensity") %>%
```

```
  activate_microbiome_dataset(what = "sample_info") %>%
  filter(sum_intensity > 20)
```

### 4.3 Merge data

Merging OTU or sample indices based on variables in the data can be a useful means of reducing noise or excess features in an analysis or graphic.

Loading included data.

```
library(microbiomedataset)
library(tidyverse)
data("global_patterns")
global_patterns
#> --------------------
#> microbiomedataset version: 0.99.1
#> --------------------
#> 1.expression_data:[ 19216 x 26 data.frame]
#> 2.sample_info:[ 26 x 8 data.frame]
#> 3.variable_info:[ 19216 x 8 data.frame]
#> 4.sample_info_note:[ 8 x 2 data.frame]
#> 5.variable_info_note:[ 8 x 2 data.frame]
#> --------------------
#> Processing information (extract_process_info())
#> create_microbiome_dataset ----------
#>             Package              Function.used                Time
#> 1 microbiomedataset create_microbiome_dataset() 2022-07-10 10:56:13
```

#### 4.4 Merge samples

Remove empty taxa.

```
global_patterns2 <-
  global_patterns %>%
  mutate2variable(what = "sum_intensity") %>%
  activate_microbiome_dataset(what = "variable_info") %>%
  dplyr::filter(sum_intensity > 0)

humantypes <- c("Feces", "Mock", "Skin", "Tongue")
global_patterns2 <-
  global_patterns2 %>%
  activate_microbiome_dataset(what = "sample_info") %>%
  dplyr::mutate(human = SampleType %in% humantypes)
```

Now on to the merging examples.

```
merged_global_patterns2 <-
  microbiomedataset::summarise_samples(object = global_patterns2,
                                       group_by = "SampleType")
extract_sample_info(merged_global_patterns2)
```

```
#>            sample_id  Primer Final_Barcode Barcode_truncated_plus_T
#> 1               Soil ILBC_01        AACGCA                   TGCGTT
#> 2              Feces ILBC_04        AAGAGA                   TCTCTT
#> 3               Skin ILBC_07        AATCGT                   ACGATT
#> 4             Tongue ILBC_10        ACACGA                   TCGTGT
#> 5         Freshwater ILBC_13        ACACTG                   CAGTGT
#> 6 Freshwater (creek) ILBC_16        ACAGCA                   TGCTGT
#> 7              Ocean ILBC_19        ACAGTT                   AACTGT
#> 8 Sediment (estuary) ILBC_22        ACATGT                   ACATGT
#> 9               Mock ILBC_27        ACCGCA                   TGCGGT
#>   Barcode_full_length         SampleType
#> 1         CTAGCGTGCGT               Soil
#> 2         TCGACATCTCT              Feces
#> 3         CGAGTCACGAT               Skin
#> 4         TGTGGCTCGTG             Tongue
#> 5         CATGAACAGTG         Freshwater
#> 6         GACCACTGCTG Freshwater (creek)
#> 7         TCGCGCAACTG              Ocean
#> 8         CACGTGACATG Sediment (estuary)
#> 9         TGACTCTGCGG               Mock
#>                                Description   class human
#> 1     Calhoun South Carolina Pine soil, pH 4.9 Subject FALSE
#> 2      M3, Day 1, fecal swab, whole body study Subject  TRUE
#> 3      M3, Day 1, right palm, whole body study Subject  TRUE
#> 4         M3, Day 1, tongue, whole body study  Subject  TRUE
#> 5 Lake Mendota Minnesota, 24 meter epilimnion  Subject FALSE
#> 6                  Allequash Creek, 0-1cm depth Subject FALSE
#> 7       Newport Pier, CA surface water, Time 1 Subject FALSE
#> 8                Tijuana River Reserve, depth 1 Subject FALSE
#> 9                                        Even1 Subject  TRUE
```

#### 4.5 Merge taxas

```
merged_variables <-
  microbiomedataset::summarize_variables(
    object = global_patterns2,
    variable_index = 1:5,
    remain_variable_info_index = 1
  )
dim(merged_variables)
#> variables   samples
#>     18984        26
dim(global_patterns2)
#> variables   samples
#>     18988        26
```
